# Enhancing Vaxign-DL for Vaccine Candidate Prediction with added ESM-Generated Features

**DOI:** 10.1101/2024.09.04.611295

**Authors:** Yichao Chen, Yuhan Zhang, Yongqun He

## Abstract

Many vaccine design programs have been developed, including our own machine learning approaches Vaxign-ML and Vaxign-DL. Using deep learning techniques, Vaxign-DL predicts bacterial protective antigens by calculating 509 biological and biomedical features from protein sequences. In this study, we first used the protein folding ESM program to calculate a set of 1,280 features from individual protein sequences, and then utilized the new set of features separately or in combination with the traditional set of 509 features to predict protective antigens. Our result showed that the usage of ESM-derived features alone was able to accurately predict vaccine antigens with a performance similar to the orginal Vaxign-DL prediction method, and the usage of the combined ESM-derived and orginal Vaxign-DL features significantly improved the prediction performance according to a set of seven scores including specificity, sensitivity, and AUROC. To further evaluate the updated methods, we conducted a Leave-One-Pathogen-Out Validation (LOPOV) study, and found that the usage of ESM-derived features significantly improved the the prediction of vaccine antigens from 10 bacterial pathogens. This research is the first reported study demonstrating the added value of protein folding features for vaccine antigen prediction.

## 1 Introduction

Vaccine design is a hot research topic, esp. after the COVID-19 pandemic. Reverse vaccinology (RV) is a method for predicting vaccine candidates by analyzing genomics and proteomics sequences [1]. There are two types of RV prediction strategies currently available. One is the filtering type, which predicts protective vaccine antigens using specific filtering criteria such as subcellular localization, adhesin probability, and the number of transmembrane helices. For example, depending on a pre-designed filtering strategy, if a protein is found to have any transmembrane helices, a vaccine antigen prediction program such as Vaxign may filter out the protein [2].The limitation of this type of method is that the evaluation criteria may not be complete and the filtering method may be too restrictive, which may lead to low sensitivity. The other strategy is machine learning based. Specifically, the second strategy is able to learn from the same set of training data and make effective predictions using specific machine learning methods such as logistic regression, support vector machine, random forest, extreme gradient boosting, and neural networks [3,4].

Our laboratory has developed many vaccine candidate prediction tools including Vaxign [2], Vaxign-ML [4], and Vaxign-DL [3]. For example, Vaxign is the first web-based reverse vaccinology method [2] Vaxign-ML [4] which is trained using a four-layer neural network to improve the prediction results. Vaxign uses the filtering method [2]Vaxign-ML uses the extreme gradient boosting method, and Vaxign-DL uses a neural network deep learning method. Vaxign is the first web-based RV method that achieved high specificity but relatively low sensitivity [2].Vaxign-ML shows high specificity and sensitivity [4].Vaxign-DL shows a performance similar to Vaxign-ML in terms of specificity and sensitivity [3].How to further improve deep learning-base vaccine antigen prediction has become a challenge.

There are many protein folding programs, including two popular deep learningbased prediction methods AlphaFold [5]and ESM (Evolutionary Scale Modeling) [6]. As an self-supervised neural language model, the AlphaFold method takes multiple sequence alignments of evolutionarily related proteins as inputs, while the ESM method requires only single input sequences. The ESM transformer protein language model scales a deep contextual language model with unsupervised learning to protein sequences with evolutionary diversity. Compared to AlphaFold, ESM runs much faster.

In this study, we hypothesized that the utilization of the intermediate features generated by a protein folding program such as the ESM program would significantly enhance our vaccine antigne prediction performance. The ESM-1b function in ESM generated 1,280 features, and our previous Vaxig-DL and VaxignML used 509 features. To evaluate our hypothesis, we calculated and used these two sets of features separately or in combination to evaluate how the usage of the ESM-generated features would enhance our vaccine antigen prediction performance. Our evaluation confirmed the value of the additional ESM-generated features in vaccine antigen prediction. In addition, we utilized our methods to predict vaccine antigens from a set of ten pathogens using a Leave-One-Pathogen-Out Validation (LOPOV) test, and further verified the value of our improved Vaxign-DL method.

## 2 Methods

### 2.1 Collection of Positive and Negative protein sequences

The training and testing data used were the same dataset used in [3] and [4]. Specifically, the dataset includes positive and negative samples. The positive samples refer to those proteins that can induce protective immune responses against virulent pathogen infections. These proteins have been verified from laboratory animal testing as valuable candidates for vaccine development. These positive sample proteins were originally downloaded from the Protegen protective antigen database [7].Additionally, since the 584 bacterial protective antigens (BPAgs) from 50 Gram-positive and Gram-negative pathogens downloaded from the Protegen database may have potential biases, homologous proteins with sequence similarity exceeding 30% have been removed in [4]. Consequently, the final positive sample data contains 397 BPAgs. To obtain negative sample data, [4]used pathogen proteins with BPAgs sequence similarity less than 30% and removed homologous proteins. These methods for obtaining positive and negative samples are consistent with those described in [8].

### 2.2 ESM generation of new features based on protein sequences

To ensure the model stability, we used the “ESM-1b” model, a pre-trained model with 33 layers and 650M parameters, and “*UR*50*/S −* 2018*−* 03” as the training set. It can eventually generate 1,280 embedding dimension features.

Because data differs between Gram positive/negative bacterial strains, the datasets of protein sequences were classified as GNN (Gram-negative protection-negative), GNP (Gram-negative protection-positive), GPN (Gram-positive protection-negative), and GPP (Gram-positive protection-positive). For example, GPP proteins are Gram-positive protection-Positive proteins that come from Gram-positive bacteria and are experimentally verified vaccine antigens. Protection-negative are negative control proteins that are likely not protective vaccine antigens. Each protein sequence data is used separately through “ESM-1b” to generate each group of results. Then, by merging the results produced by GNP and GPP, we can get the merged positive samples. Similarly, we can also get a combined protection-negative data by merging GNN and GPN feature results. Eventually We got two datasets, protection Positive and Negative. Finally, by combining the results generated by ESM-1b with our original Vaxign-ML and Vaxign-DL data [3,4],we got two sets of combined datasets with 1,789 features each.

Note that the ESM-1b model can only use up to 1,024 amino acids for the ESM protein folding prediction process. For those large size proteins, the additional amino acids beyond the 1,024 amino acids cannot be used for ESM calculation. Our study found only a small number of proteins in our datasets have the size over 1,024 amino acids. To deal with this situation, two methods were applied. The first method was to skip those proteins that have the protein sequence length exceeding 1,024 amino acids. The final data obtained is 372 positive samples and 3,969 negative sample, both have 1,789 features. Therefore, our second method was to use only the first 1,024 amino acids if a protein has over 1,024 amino acids. The results of the two methods were also compared and analyzed.

### 2.3 Leave-one-pathogen-out validation

To test vaccine candidates against novel pathogens, we retained the [3] Leave-One-Pathogen-Out Validation (LOPOV) dataset from the Vaxign-ML and Vaxign-DL studied, and calculated and added 1,280 features generated by ESM to the original datasets. This approach involved a set of ten pathogens, including four Gram-positive pathogens (*Mycobacterium tuberculosis, Staphylococcus aureus, Streptococcus pneumoniae*, and *Streptococcus pyogenes*) and six Gramnegative pathogens (*Helicobacter pylori, Neisseria meningitidis, Brucella* spp., *Escherichia coli, Yersinia pestis*, and *Hemophilus influenzae*). For each LOPOV analysis, the positive and negative protein samples from one specific pathogen were left out, and the samples from the remaining nine pathogens were used for model training. The trained model with the nine pathogen samples was then used to predict vaccine antigens from the left-out pathogen.

### 2.4 Deep learning pipeline

The overall pipeline is shown in Fig. 1. Through the ESM protein folding program, we got 1,280 features generated by the ESM pre-trained model. Different from the biological or chemical features from the traditional Vaxign-ML method, these ESM features are named “Feature 1”, “Feature 2”, etc, so it is difficult to identify the specific meaning of these ESM features. By combine these new ESM features with 509 features from the original Vaxign-DL/Vaxign-ML studies, we obtained the total of 1,789 features for each protein.

**Fig. 1.**
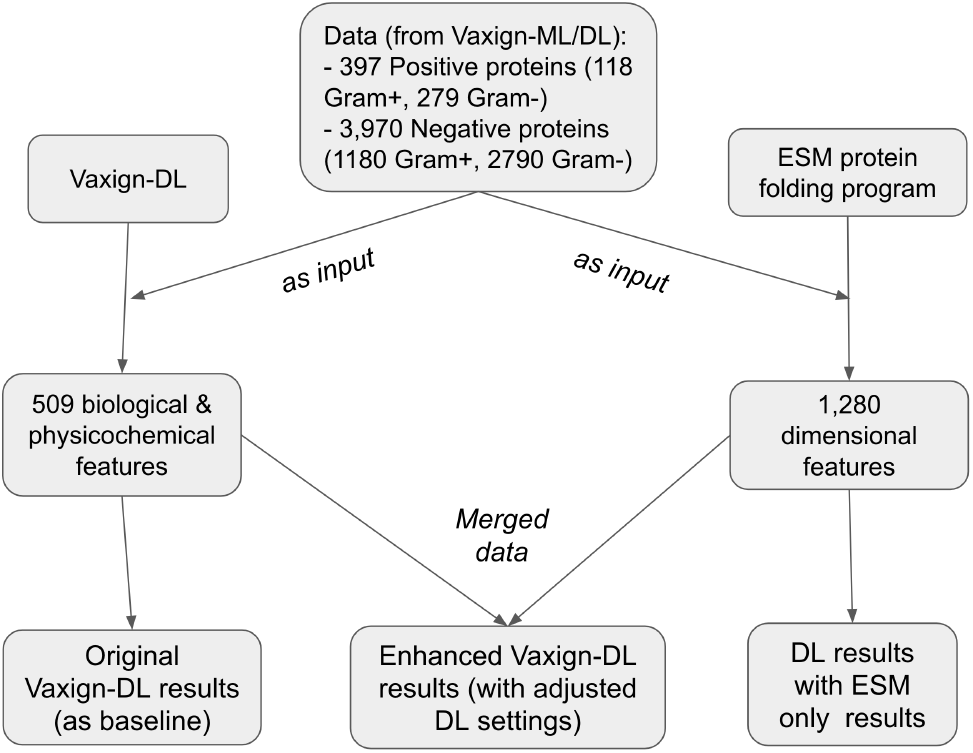
Overall Pipeline

The original data samples include 397 positive protein samples and 3,970 negative protein samples. After we removed those protein sequences that have over 1,024 amino acids, our data samples used for ESM processing are only 372 positive samples. To maintain the original 1:10 data ratio, we randomly deleted many negative samples, resulting in the number of 3,720 negative protein samples. Due to the change of the sample sizes, we fine-tuned the weights of the data in our deep learning process.

Our deep learning method used Multi-layer Perceptrons (MLP) model, the same classic deep learning model as used in Vaxign-DL [3]. The MLP model uses the sequential layering of nonlinear processing units and the back-propagation optimization method [9].In addition to the same three-layer hidden MLP model as used in Vaxign-DL, we also tested a five-layer hidden MLP model in our study. The two middle hidden layers have 256 units and 128 units respectively. Each fully connected layer used LeakyReLU [10]activation function with a slight slope of 0.005 to maintain activity. After LeakyReLU, batch normalization was applied to improve optimization and stability. In addition, in order to increase Robustness and prevent Overfitting, each fully connected layer also had dropout layers with a dropout rate of 0.1. The final output layer used softmax activation function to ensure that the result is the probability of each category.

### 2.5 Performance evaluation

For the evaluation model, in order to be able to clearly compare the results, the original evaluation criteria were applied. Specifically, our result evaluation calculated seven scores: value accuracy, sensitivity, specificity, weighted F1 score, Matthews correlation coefficient (MCC), Area under the Receiver Operating Characteristic (AUROC), and Area under Precision-Recall Curve (AUPRC).

## 3 Results

### 3.1 Performance analysis ESM enhance Vaxign-DL model

Using the 372 protection-positive and 3,720 protection-negative samples, we tested and compared the prediction performances with three different sets of features: the original Vexing-DL set of 509 features, only ESM-generated 1,280 features, and the set of 1,789 features by combining the original set and the new ESM features (Table 1). By calculating the seven evaluation scores as defined in the Methods, we found that overall original Vaxign-DL method (baseline) and the ESM only method achieved similar prediction performances and the method with combined features method achieved higher performances in all scores expect that the three methods all achieve the same high specificity score of 0.99 + 0.004.

**Table 1.**
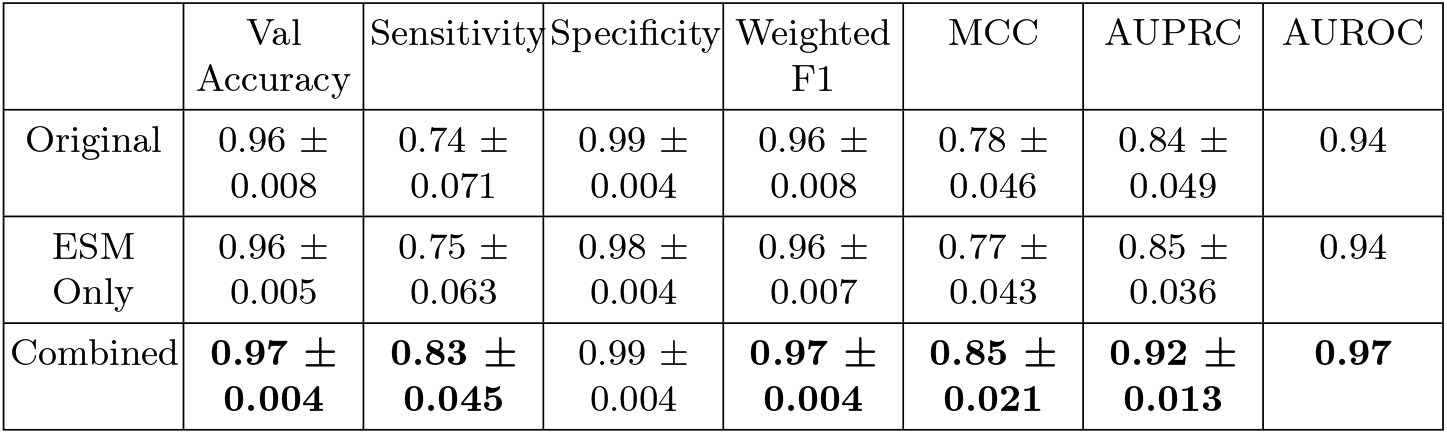
Performance metrics for different models.

Out of the seven scores calculated, all except sensitivity and MCC were over 0.9 with all the three methods, suggesting the high performances for all the methods. The combined method achieved achieved a value accuracy of 0.97, sensitivity of 0.83, wighted F1 score of 0.97, MCC of 0.85, AUPRC of 0.92, and AUROC of 0.97, which were all higher than the scored generated from the original and ESM only methods. For example, the original, ESM only, and combined method achieved an AUROC score of 0.94, 0.94, and 0.97, respectively; and they achieved a sensitivity score of 0.74, 0.75, and 0.83, respectively. Given the specificity scores similar among the three methods, the higher sensitivity score from the combined method might be an important factor to its higher AUROC score.

Overall, additional features from the ESM protein folding program helped the Vaxign-DL model achieve better performance of protective vaccine antigen prediction.

### 3.2 Hyperparameter Optimization Study

In the Vexign-DL model, four hidden layers were originally used [3]. In order to further improve the prediction results of Vaxign-DL combined with ESM, we further explored whether more Neural Network layers could improve the prediction results of the model. To this end, we added a fully connected layer to the original model, 256, 128, 64. After testing the five-layer fully connected layer, it was found that more layers did not significantly improve the performance (Fig. 2). For this reason, we changed the dimensions of the fully connected layer to 512, 256, 128. The results were consistent and there was no significant change between these two settings. The four-layer model indeed achieved the best sensitivity (0.83 ± 0.045) (Table 2). In the end, the original four-layer fully connected layer model was retained as our optimal setting in the later studies.

**Table 2.**
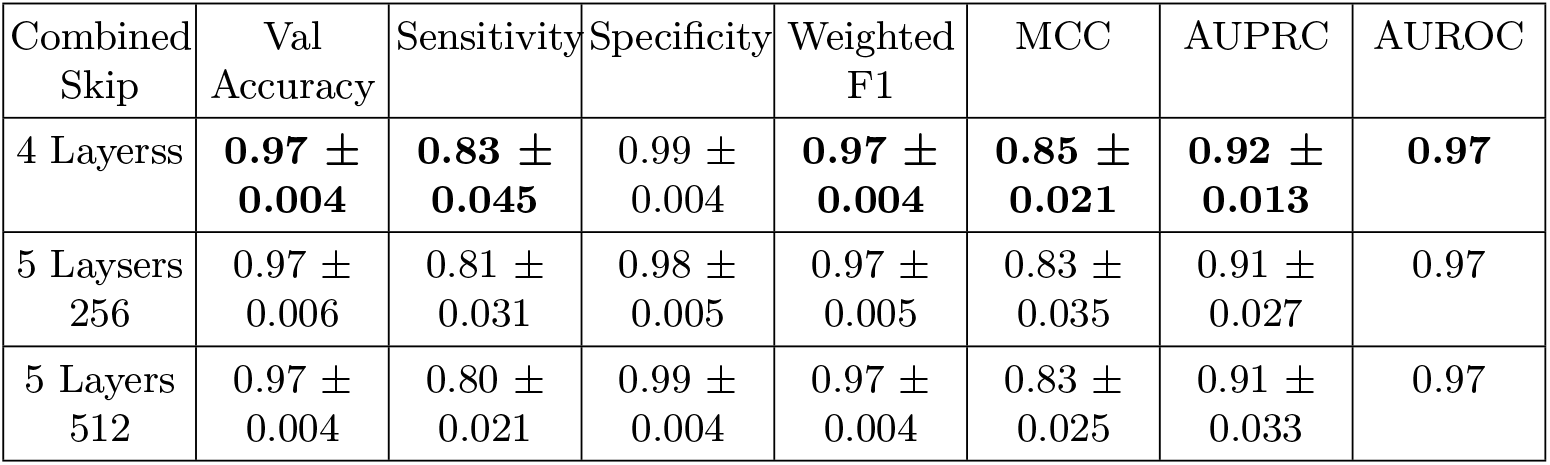
Performance metrics for different number of Layers.

| Combined Skip | Val Accuracy | Sensitivity | Specificity | Weighted F1 | MCC | AUPRC | AUROC |
| --- | --- | --- | --- | --- | --- | --- | --- |
| 4 Layers | **0.97 ± 0.004** | **0.83 ± 0.045** | 0.99 ± 0.004 | **0.97 ± 0.004** | **0.85 ± 0.021** | **0.92 ± 0.013** | **0.97** |
| 5 Layers 256 | 0.97 ± 0.006 | 0.81 ± 0.031 | 0.98 ± 0.005 | 0.97 ± 0.005 | 0.83 ± 0.035 | 0.91 ± 0.027 | 0.97 |
| 5 Layers 512 | 0.97 ± 0.004 | 0.80 ± 0.021 | 0.99 ± 0.004 | 0.97 ± 0.004 | 0.83 ± 0.025 | 0.91 ± 0.033 | 0.97 |

**Fig. 2.**
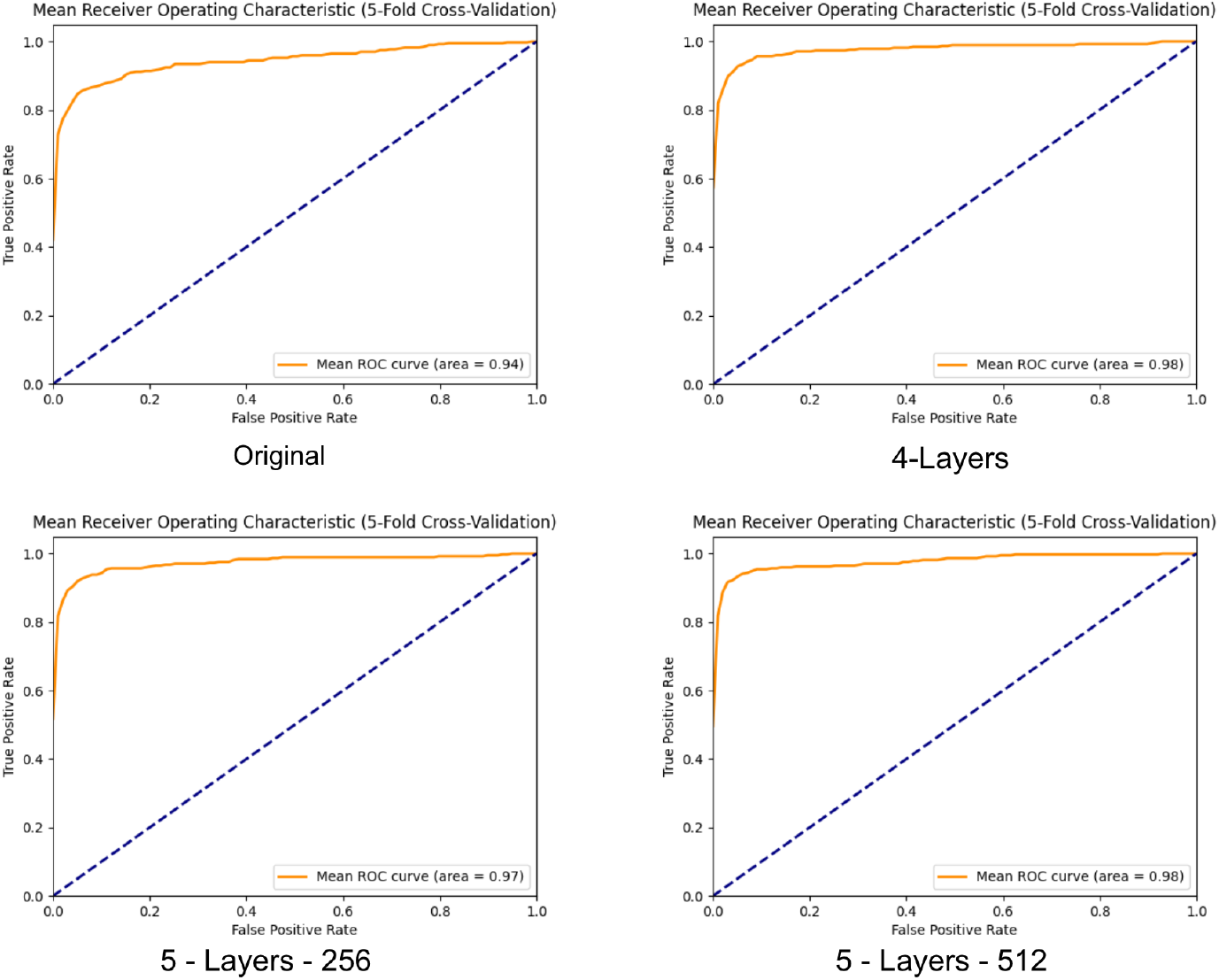
Mean ROC results

#### Comparison of Two ESM methods for Processing Long Sequence Proteins

Out of the 397 positive vaccine antigens in our dataset, 25 (6%) have the size over 1,024 amino acids. Since ESM can generate features for proteins up to 1,024 amino acids, we designed and tested two different methods to include these large size proteins. The “Skip” way was that when the amino acid length of a protein is over 1,024, the protein would be skipped for further usage in prediction. The “Cut” method was that when the amino acid length is over 1,024, only the first 1,024 amino acids were used as input. Table 3 shows the results of the two different methods. The “Skip” method achieved better results in all aspects. It is possible that the “Cut” could not get the accurate prediction of all the features for the large-size protein sequence using ESM-1b, which might have led to a bias in the prediction.

**Table 3.** Performance metrics for different ESM Processing.

| ESM-1b | Val Accuracy | Sensitivity | Specificity | Weighted F1 | MCC | AUPRC | AUROC |
| --- | --- | --- | --- | --- | --- | --- | --- |
| Skip | **0.97 ± 0.004** | **0.83 ± 0.045** | **0.99 ± 0.004** | **0.97 ± 0.004** | **0.85 ± 0.021** | **0.92 ± 0.013** | **0.97** |
| Cut | 0.96 ± 0.005 | 0.75 ± 0.063 | 0.98 ± 0.004 | 0.96 ± 0.007 | 0.77 ± 0.043 | 0.85 ± 0.036 | 0.94 |

#### Leave-one-pathogen-out Validation

To further evaluate different deep learning methods tested in this study, we applied the Leave-one-pathogen-out Validation (LOPOV) approach, which tested how a deep learning method learned from using only 9 bacterial pathogens’ data could be used to predict vaccine candidates from another new bacterial pathogen. This was the method originally used in Vaxign-ML [4].The scores, the Area under the Receiver Operating Characteristic (ROC) score and the Area under the Precision-Recall curve (ROC) score were calculated (Fig. 3). Our results showed that the “Skip” and “Cut” methods that used the ESM-derived features in combination with the original Vaxign-DL features both performed better than the original Vaxign-DL method. The Cut method achieved an average AUC of 0.98 and AUPRC of 0.94, and Skip method achieved an average AUC of 0.98 and AUPRC of 0.93, both were better than the Original method results of AUC of 0.95 and AUPRC of 0.85. These confirmed the improved performance using the ESM pre-training model.

**Fig. 3.**
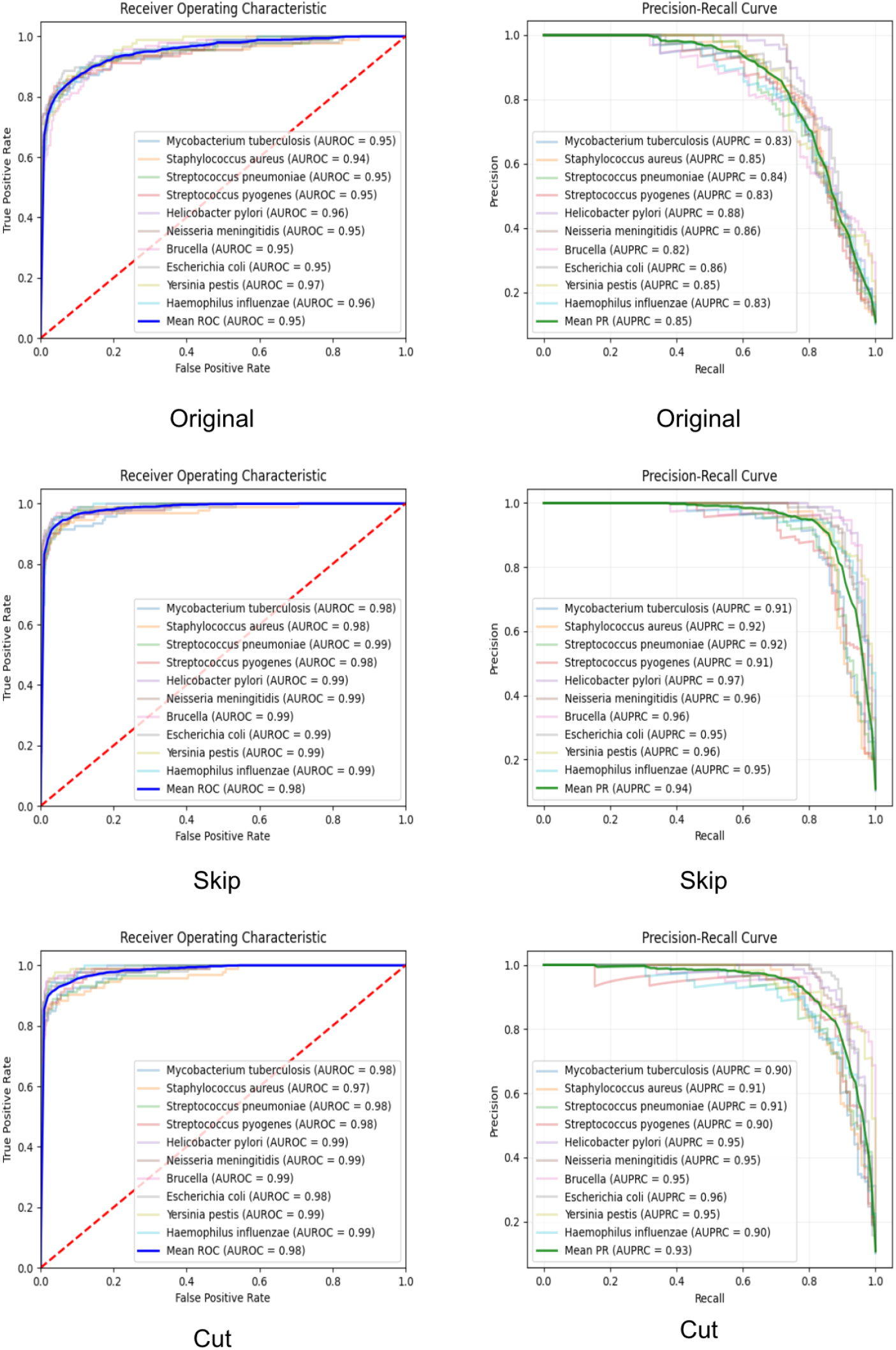
LOPOV evaluations using different deep learning methods

## 4 Discussion

To our best knowledge, our work represents the first deep learning-based vaccine antigen prediction program that uses the intermediate features out of a protein folding program (i.e., ESM). Our results showed that such a protein folding based method enhanced the vaccine prediction performance. The improved performance of the enhanced Vaxign-DL model for vaccine antigen prediction using features generated from a protein folding prediction program has significant implications and applications for accelerating vaccine development processes.

The authors’ consideration of future work demonstrates a forward-thinking approach. The potential use of AlphaFold to address the protein length limitation is particularly interesting to me. In conclusion, while there are areas for improvement, the scientific merit and novelty of the work are clear.

In this study, the ESM protein folding program was used due to its advantage of fast run time and accuracy. It is also possible to use other protein folding methods for vaccine antigen prediction, such as AlphaFold. One primary reason we did not use AlphaFold is that AlphaFold runs much slower than the ESM method. However, given the other advantages of AlphaFold, we plan to test the usage of AlphaFold in our further study. In addition, we will also evaluate the statistical significance of improvements among different methods.

We note that the abstract nature of the 1280 ESM-derived features may pose challenges to the interpretability of the model. The introduction of these features is intended to enhance the predictive power of the model, which is critical in vaccine development. However, represented as high-dimension embeddings, these features appear to be unlikely linked to specific biological meanings. We recognize that in this field, it is equally important to understand the basis of model predictions. In future work, we will explore in more depth how to balance the predictive performance and interpretability of the model, identify possible biological relevance of specific features, and consider introducing more methods of interpretability analysis to better support the needs of vaccine development.

One disadvantage of the ESM program is that it can only calculate features for proteins with the maximum of 1,024 amino acids. Bacterial proteins have the average size of 320 amino acids [11]. In our study, 25 out of 397 (6%) positive vaccine antigens have the size over 1,024 amino acids, indicating that the number of oversized proteins is small. We tried the “Skip” and “Cut” methods to deal with this situation. More solutions can be tried later, such as sliding window and subsequence selection. The sliding window technology extracts multiple overlapping fragments from long sequences for embedding, and then aggregates the embedding results of these fragments. The subsequence selection method selects important subsequences in the long sequences for embedding and ignores unimportant parts. In future studies, we plan to further analyze the characteristics of these long proteins, including whether they have common features, whether they represent specific antigenic classes, and their distributions in different bacteria. These will help us more fully understand the impact of these proteins on model performance and potentially develop better methods to handle this type of protein.

More other experiments can be explored in the future. In our ESM pre-trained model study, we conservatively only used the “ESM-1b” model. There have been many “ESM2” models available for testing, such as the “ESM2-t48-15B-UR50D” model [12], which has 15 billion parameters, a 48-layer Transformer architecture, and multi-task learning [13].These models will require more computational power and may generate further enhanced results. In addition, our DL model currently only uses the multilayer perceptron (MLP) method. We may evaluate the performance of other methods such as Long Short-Term Memory (LTSM) [14]and use more complex DL networks to strengthen the model.

## Acknowledgments

This research is funded by a NIH-NIAID grant U24AI171008. We also acknowledge the usage of the training data originally generated in the published Vaxign-ML study.

